## Supplementary Information for "Holimap: an accurate and efficient method for solving stochastic gene network dynamics"

### Supplemental Material —
### Holimap: an accurate and efficient method for solving stochastic gene networks

Chen Jia, Ramon Grima

### Note S1. Technical details of figures

#### Technical details for Fig. 2

In (e), the parameters are chosen as $h = 1, d = 1, B = 0.1, \sigma_u = 2, \sigma_b = 0.2$.

In (f), the parameters are $h = 2, d = 1, B = 0.1, \rho_u = 5, \rho_b = 200$.

In (g), the parameters are as in (f). The unbinding and binding rates are chosen as $\sigma_u = 0.01, \sigma_b = 0.015$ for slow gene switching, $\sigma_u = 1, \sigma_b = 0.06$ for intermediate fast gene switching, and $\sigma_u = 100, \sigma_b = 1.8$ for fast gene switching.

In (h), the parameters are $d = 1, B = 0.1, \rho_u = 50, \rho_b = 200$. The maximum HD is computed when $\sigma_u$ varies from $10^{-2}$ to $10^2$, $\sigma_b$ varies from $10^{-2-h}$ to $10^{2-h}$ and the other parameters remain fixed.

#### Technical details for Fig. 3

In (d), the parameters are chosen as $h_1 = h_2 = 1, d_1 = d_2 = 1, \rho_{u1} = 21, \rho_{b1} = 7, \rho_{u2} = 0, \rho_{b2} = 3, \sigma_{u1} = \sigma_{u2} = 5$.

In (e), the parameters are as in (d). The binding rates are chosen as $\sigma_{b1} = 10^3, \sigma_{b2} = 4 \times 10^{-4}$.

In (g), the parameters are $h_1 = h_2 = 2, d_1 = d_2 = 1, \rho_{u1} = 18, \rho_{b1} = 1, \rho_{u2} = 2, \rho_{b2} = 0, \sigma_{u1} = \sigma_{u2} = 5$.

In (h), the parameters are as in (g). The binding rates are chosen as $\sigma_{b1} = 10^3, \sigma_{b2} = 1.5$.

#### Technical details for Fig. 4

In (c),(d), the parameters are chosen as $h_1 = h_2 = h_3 = 3, d_1 = d_2 = d_3 = 1, \rho_{u1} = \rho_{u2} = \rho_{u3} = 81, \rho_{b1} = 5.4, \rho_{b2} = \rho_{b3} = 0, \sigma_{u1} = \sigma_{u2} = \sigma_{u3} = 1.1, \sigma_{b1} = \sigma_{b2} = 0.04, \sigma_{b3} = 4$. The time-dependent protein distributions for the LMA are computed using the time-average method (a method of determination the effective parameter) described in Ref. [1], while those for the 2-HM are computed by applying FSP directly to the CME of the linear network with time-dependent rates (upper right panel of Fig. 4(a) in the main text).

#### Technical details for Fig. 6

In (g),(h), there are four gene networks whose parameter values are described as follows.

For the network with autoregulation and protein sequestration, the parameters are chosen as $h_1 = h_2 = 1, d_1 = d_2 = 1, \alpha = 0.3, \rho_{u1} = 13.5, \rho_{u2} = 0, \rho_{b1} = \rho_{b2} = 81, \sigma_{u1} = \sigma_{u2} = 0.8, \sigma_{b1} = 0.024, \sigma_{b2} = 0.033$.

For the network with autoregulation and protein phosphorylation, the parameters are chosen as $h = 1, d = 1, d_1 = 0.1, d_2 = 3, \rho_u = 30, \rho_b = 150, \sigma_u = 1, \sigma_b = 0.17, a_1 = a_3 = b_2 = b_4 = c_1 = c_3 = 1, a_2 = a_4 = b_1 = b_3 = c_2 = c_4 = 0.1$. The total number of each enzyme (including the free enzyme $E_i$ and the complex $C_i$) is chosen as 100.

For the network with autoregulation and mRNA degradation control, the parameters are chosen as $h = 1, v = 1, d = 0.2, \rho_u = 105, \rho_b = 21, \sigma_u = \sigma_b = 2, u = 3, \alpha = 0.2, a = b = 1$. The total number of enzyme (including the inactive form $E$ and the active form $E^*$) is chosen as 100.

For the network with microRNA-mRNA interactions, the parameters are chosen as $h_1 = h_2 = 1, d_1 = d_2 = 1, \alpha = 0.3, \rho_{u1} = 15, \rho_{u2} = 0, \rho_{b1} = \rho_{b2} = 90, \sigma_{u1} = \sigma_{u2} = \sigma_{b1} = 1, \sigma_{b2} = 0.033, \alpha = 0.1, \beta = 0.001, a_1 = b_1 = 2, a_2 = b_2 = 0.5$.

#### Technical details for Supplementary Fig. S6

In (c), there are two types of network topologies. For the network in the left panel, the parameters are chosen as $d_1 = d_2 = 1, \rho_H = \rho_{uu1} = 28, \rho_L = \rho_{bu1} = \rho_{ub1} = \rho_{bb1} = 7, \rho_{u2} = 20, \rho_{b2} = 0, K = 100, L = 0.1, \lambda_{u1} = \sigma_{u1} = 0.9, \lambda'_{u1} = \sigma'_{u1} = 100, \lambda_{b1} = \lambda_{u1}/K, \sigma_{b1} = \sigma_{u1}/L, \lambda'_{b1} = \lambda'_{u1}L/M, \sigma'_{b1} = \sigma'_{u1}K/M, \sigma_{u2} = 1, \sigma_{b2} = 7,$

where the parameter $M$ is chosen as $M = KL$ for independent binding, $M = 10^{-4}KL$ for positive cooperative binding, and $M = 10^4 KL$ for negative cooperative binding.

For the network in the right panel, the parameters are chosen as $d_1 = d_2 = 1, \rho_H = \rho_{bu1} = 28, \rho_L = \rho_{uu1} = \rho_{ub1} = \rho_{bb1} = 7, \rho_{u2} = 20, \rho_{b2} = 0, K = 0.1, L = 0.1, \lambda_{u1} = \sigma_{u1} = 10, \lambda'_{u1} = \sigma'_{u1} = 100, \lambda_{b1} = \lambda_{u1}/K, \sigma_{b1} = \sigma_{u1}/L, \lambda'_{b1} = \lambda'_{u1}L/M, \sigma'_{b1} = \sigma'_{u1}K/M, \sigma_{u2} = 1, \sigma_{b2} = 7$, where the parameter $M$ is chosen as $M = KL$ for independent binding, $M = KL/2$ for positive cooperative binding, and $M = 10KL$ for negative cooperative binding.

In (d), we randomly select $10^4$ sets of parameters for each network topology and each type of transcription factor interaction. Specifically, the parameters are chosen as follows:

• $d_1 = d_2 = 1$,

• $\rho_H \sim U[10, 50], \rho_L \sim U[0, 10], \max\{\rho_{u2}, \rho_{b2}\} \sim U[2, 10], \min\{\rho_{u2}, \rho_{b2}\} \sim U[0, 2]$,

• $\log_{10} K, \log_{10} L \sim U[-2, 2]$, $M = KL$ for independent binding, $\log_{10}(M/KL) \sim U[-3, 0]$ for positive cooperative binding, and $\log_{10}(M/KL) \sim U[0, 3]$ for negative cooperative binding,

• $\lambda_{u1}, \sigma_{u1}, \lambda'_{u1}, \sigma'_{u1}, \sigma_{u2}, \sigma_{b2} \sim U[0.1, 10]$,

• $\lambda_{b1} = \lambda_{u1}/K, \sigma_{b1} = \sigma_{u1}/L, \lambda'_{b1} = \lambda'_{u1}L/M, \sigma'_{b1} = \sigma'_{u1}K/M$,

where $U[x, y]$ denotes the uniform distribution over the interval $[x, y]$, $\max\{x, y\}$ denotes the larger one between $x$ and $y$, and $\min\{x, y\}$ denotes the smaller one between $x$ and $y$.

### Note S2. Holimaps for autoregulatory feedback loops

We consider an autoregulatory feedback loop with bursty protein expression and cooperative protein binding; this is illustrated in Fig. 2(a) in the main text. It is described by the following reactions:

$$G + hP \xrightarrow{\sigma_b} G^*, \quad G^* \xrightarrow{\sigma_u} G + hP,$$
$$G \xrightarrow{\rho_u} G + kP, \quad G^* \xrightarrow{\rho_b} G^* + kP, \quad P \xrightarrow{d} \varnothing,$$

where $\sigma_b$ is the binding rate of protein to the gene; $\sigma_u$ is the unbinding rate; $\rho_u$ and $\rho_b$ are the burst frequencies when gene is in the unbound and bound states, respectively; $d$ is the degradation rate of protein; $k$ is the burst size of protein which is geometrically distributed with parameter $p$. Feedback is mediated by cooperative binding of $h$ protein copies to the gene. The reaction scheme describes a positive feedback loop when $\rho_b > \rho_u$ and describes a negative feedback loop when $\rho_b < \rho_u$.

The microstate of the gene of interest can be represented by an ordered pair $(i, n)$, where $n$ is the number of protein molecules and $i$ is the state of the gene with $i = 0, 1$ corresponding to the unbound and bound states, respectively. Let $p_{i,n}$ denote the probability of having $n$ protein molecules when the gene is in state $i$. Then the dynamics of the system is governed by the CMEs

$$\dot{p}_{0,n} = \left[\sum_{k=0}^{n-1} \rho_u p^{n-k} q p_{0,k} - \sum_{k=1}^{\infty} \rho_u p^k q p_{0,n}\right] + d[(n+1)p_{0,n+1} - np_{0,n}] + \left[\sigma_u p_{1,n-h} - \sigma_b \frac{n!}{(n-h)!} p_{0,n}\right],$$
$$\dot{p}_{1,n} = \left[\sum_{k=0}^{n-1} \rho_b p^{n-k} q p_{1,k} - \sum_{k=1}^{\infty} \rho_b p^k q p_{1,n}\right] + d[(n+1)p_{1,n+1} - np_{1,n}] + \left[\sigma_b \frac{(n+h)!}{n!} p_{0,n+h} - \sigma_u p_{1,n}\right],$$

where $q = 1 - p$. Here the first term on the right-hand side describes protein synthesis, the second term describes protein degradation, and the third term describes gene state switching. We next define the generating function

$$F_i(x) = \sum_{n=0}^{\infty} p_{i,n} x^n, \quad i = 0, 1.$$

Then the CMEs can be converted into the PDEs

$$\begin{aligned}\partial_t F_0 &= \frac{\rho_u p(x-1)}{1-px}F_0 - d(x-1)\partial_x F_0 + \sigma_u x^h F_1 - \sigma_b x^h \partial_x^{(h)} F_0,\\ \partial_t F_1 &= \frac{\rho_b p(x-1)}{1-px}F_1 - d(x-1)\partial_x F_1 - \sigma_u F_1 + \sigma_b \partial_x^{(h)} F_0.\end{aligned} \tag{1}$$

To proceed, let

$$g_i = \sum_{n=0}^{\infty} p_{n,i} = F_i(1) \tag{2}$$

be the probability that the gene is in state $i$ and let

$$\mu_{k,i} = \sum_{n=0}^{\infty} n(n-1)\cdots(n-k+1)p_{n,i} = \partial_x^{(k)} F_i(1) := \frac{d^k}{dx^k}\bigg|_{x=1} F_i(x) \tag{3}$$

be the $k$th factorial moment of the protein number when the gene is in state $i$. Combining Eqs. (1), (2), and (3), it is easy to check that the evolution of zero, first, and second-order moments is given by

$$\begin{aligned}\dot{g}_0 &= \sigma_u g_1 - \sigma_b \mu_{h,0},\\ \dot{\mu}_{1,0} &= \rho_u B g_0 - d\mu_{1,0} + \sigma_u(\mu_{1,1} + hg_1) - \sigma_b(\mu_{h+1,0} + h\mu_{h,0}),\\ \dot{\mu}_{1,1} &= \rho_b B g_1 - d\mu_{1,1} - \sigma_u \mu_{1,1} + \sigma_b \mu_{h+1,0},\\ \dot{\mu}_{2,0} &= 2\rho_u B(\mu_{1,0} + Bg_0) - 2d\mu_{2,0} + \sigma_u(\mu_{2,1} + 2h\mu_{1,1} + h(h-1)g_1)\\ &\quad - \sigma_b(\mu_{h+2,0} + 2h\mu_{h+1,0} + h(h-1)\mu_{h,0}),\\ \dot{\mu}_{2,1} &= 2\rho_b B(\mu_{1,1} + Bg_1) - 2d\mu_{2,1} - \sigma_u \mu_{2,1} + \sigma_b \mu_{h+2,0},\end{aligned} \tag{4}$$

where $g_1 = 1 - g_0$ and $B = \sum_{n=0}^{\infty} np^n(1-p) = p/(1-p)$ is the mean burst size of protein.

#### Algorithm for the 2-HM

We next consider the 2-HM, which maps the nonlinear network to the following linear one:

$$G \xrightarrow{\tilde{\sigma}_b} G^*, \quad G^* \xrightarrow{\tilde{\sigma}_u} G,$$
$$G \xrightarrow{\rho_u} G + kP, \quad G^* \xrightarrow{\rho_b} G^* + kP, \quad P \xrightarrow{d} \varnothing,$$

where $\tilde{\sigma}_b$ and $\tilde{\sigma}_u$ are the effective rates of gene state switching. Next we determine the values of the two effective parameters. For the linear network, the generating functions satisfy the PDEs

$$\begin{aligned}\partial_t F_0 &= \frac{\rho_u p(x-1)}{1-px}F_0 - d(x-1)\partial_x F_0 + \sigma_u F_1 - \sigma_b F_0,\\ \partial_t F_1 &= \frac{\rho_b p(x-1)}{1-px}F_1 - d(x-1)\partial_x F_1 - \sigma_u F_1 + \sigma_b F_0.\end{aligned} \tag{5}$$

Combining Eqs. (2), (3), and (5), we find that the evolution of zeroth, first, and second-order moments are given by

$$\begin{aligned}\dot{g}_0 &= \tilde{\sigma}_u g_1 - \tilde{\sigma}_b g_0,\\ \dot{\mu}_{1,0} &= \rho_u B g_0 - d\mu_{1,0} + \tilde{\sigma}_u \mu_{1,1} - \tilde{\sigma}_b \mu_{1,0},\\ \dot{\mu}_{1,1} &= \rho_b B g_1 - d\mu_{1,1} - \tilde{\sigma}_u \mu_{1,1} + \tilde{\sigma}_b \mu_{1,0},\\ \dot{\mu}_{2,0} &= 2\rho_u B(\mu_{1,0} + Bg_0) - 2d\mu_{2,0} + \tilde{\sigma}_u \mu_{2,1} - \tilde{\sigma}_b \mu_{2,0},\\ \dot{\mu}_{2,1} &= 2\rho_b B(\mu_{1,1} + Bg_1) - 2d\mu_{2,1} - \tilde{\sigma}_u \mu_{2,1} + \tilde{\sigma}_b \mu_{2,0}.\end{aligned} \tag{6}$$

The two effective parameters $\tilde{\sigma}_u$ and $\tilde{\sigma}_b$ are chosen so that the two systems have the same zeroth and first-order moment equations (for the latter, we mean the first-order moment when the gene is in the bound state). Matching the first and third identities in Eqs. (4) and (6), we find that $\tilde{\sigma}_b$ and $\tilde{\sigma}_u$ should be determined by the following system of linear equations:

$$\begin{aligned}\tilde{\sigma}_u g_1 - \tilde{\sigma}_b g_0 &= \sigma_u g_1 - \sigma_b \mu_{h,0},\\ \tilde{\sigma}_u \mu_{1,1} - \tilde{\sigma}_b \mu_{1,0} &= \sigma_u \mu_{1,1} - \sigma_b \mu_{h+1,0}.\end{aligned} \tag{7}$$

Solving these equations yields

$$\begin{aligned}\tilde{\sigma}_u &= \sigma_u - \sigma_b \frac{g_0\mu_{h+1,0} - \mu_{h,0}\mu_{1,0}}{g_0\mu_{1,1} - g_1\mu_{1,0}},\\ \tilde{\sigma}_b &= \sigma_b \frac{\mu_{h,0}\mu_{1,1} - g_1\mu_{h+1,0}}{g_0\mu_{1,1} - g_1\mu_{1,0}}.\end{aligned} \tag{8}$$

In the case of non-cooperative binding ($h = 1$), we note that $\tilde{\sigma}_u = \tilde{\sigma}_u(g_i, \mu_{1,i}, \mu_{2,i})$ and $\tilde{\sigma}_b = \tilde{\sigma}_b(g_i, \mu_{1,i}, \mu_{2,i})$ are functions of $g_i$, $\mu_{1,i}$, and $\mu_{2,i}$, and hence inserting Eq. (8) into Eq. (6) yields a set of closed moment equations, i.e.

$$\begin{aligned}\dot{g}_0 &= \tilde{\sigma}_u(g_i, \mu_{1,i}, \mu_{2,i})g_1 - \tilde{\sigma}_b(g_i, \mu_{1,i}, \mu_{2,i})g_0,\\ \dot{\mu}_{1,0} &= \rho_u B g_0 - d\mu_{1,0} + \tilde{\sigma}_u(g_i, \mu_{1,i}, \mu_{2,i})\mu_{1,1} - \tilde{\sigma}_b(g_i, \mu_{1,i}, \mu_{2,i})\mu_{1,0},\\ \dot{\mu}_{1,1} &= \rho_b B g_1 - d\mu_{1,1} - \tilde{\sigma}_u(g_i, \mu_{1,i}, \mu_{2,i})\mu_{1,1} + \tilde{\sigma}_b(g_i, \mu_{1,i}, \mu_{2,i})\mu_{1,0},\\ \dot{\mu}_{2,0} &= 2\rho_u B(\mu_{1,0} + B g_0) - 2d\mu_{2,0} + \tilde{\sigma}_u(g_i, \mu_{1,i}, \mu_{2,i})\mu_{2,1} - \tilde{\sigma}_b(g_i, \mu_{1,i}, \mu_{2,i})\mu_{2,0},\\ \dot{\mu}_{2,1} &= 2\rho_b B(\mu_{1,1} + B g_1) - 2d\mu_{2,1} - \tilde{\sigma}_u(g_i, \mu_{1,i}, \mu_{2,i})\mu_{2,1} + \tilde{\sigma}_b(g_i, \mu_{1,i}, \mu_{2,i})\mu_{2,0}.\end{aligned}$$

where $g_1 = 1 - g_0$. Solving the above equations, we can obtain the approximate values of all zeroth, first, and second-order moments ($g_i$, $\mu_{1,i}$, and $\mu_{2,i}$). Finally we can use Eq. (8) to determine the values of $\tilde{\sigma}_b$ and $\tilde{\sigma}_u$.

We emphasize that moment closure can also be done alternatively as follows. When $h = 1$, combining the first three identities in Eq. (4) and the last two identities in Eq. (6), and then inserting Eq. (8) into Eq. (6) also yield a set of closed moment equations, i.e.

$$\begin{aligned}\dot{g}_0 &= \sigma_u g_1 - \sigma_b \mu_{1,0},\\ \dot{\mu}_{1,0} &= \rho_u B g_0 - d\mu_{1,0} + \sigma_u(\mu_{1,1} + g_1) - \sigma_b(\mu_{2,0} + \mu_{1,0}),\\ \dot{\mu}_{1,1} &= \rho_b B g_1 - d\mu_{1,1} - \sigma_u\mu_{1,1} + \sigma_b\mu_{2,0},\\ \dot{\mu}_{2,0} &= 2\rho_u B(\mu_{1,0} + B g_0) - 2d\mu_{2,0} + \tilde{\sigma}_u(g_i, \mu_{1,i}, \mu_{2,i})\mu_{2,1} - \tilde{\sigma}_b(g_i, \mu_{1,i}, \mu_{2,i})\mu_{2,0},\\ \dot{\mu}_{2,1} &= 2\rho_b B(\mu_{1,1} + B g_1) - 2d\mu_{2,1} - \tilde{\sigma}_u(g_i, \mu_{1,i}, \mu_{2,i})\mu_{2,1} + \tilde{\sigma}_b(g_i, \mu_{1,i}, \mu_{2,i})\mu_{2,0}.\end{aligned}$$

We find that the above two closure methods lead to very similar results. Our moment closure method essentially assumes that the marginal dynamics of a complex nonlinear network can be approximated by that of a decoupled linear network and it provides a way to find the optimal approximation. The advantage of our moment closure method is that it does not assume in advance the shape of the protein distribution; specifically, it does not assume the protein distribution to be of a simple type such as the Gaussian, Poisson, Log-normal, or Gamma distributions, as commonly assumed by conventional moment closure methods [2–5].

In the case of cooperative binding ($h \geq 2$), we need higher-order moment equations. For example, direct computations show that the evolution of third and fourth moments are given by

$$\begin{aligned}\dot{\mu}_{3,0} &= 3\rho_u B(\mu_{2,0} + 2B\mu_{1,0} + 2B^2 g_0) - 3d\mu_{3,0} + \tilde{\sigma}_u\mu_{3,1} - \tilde{\sigma}_b\mu_{3,0},\\ \dot{\mu}_{3,1} &= 3\rho_b B(\mu_{2,1} + 2B\mu_{1,1} + 2B^2 g_1) - 3d\mu_{3,1} - \tilde{\sigma}_u\mu_{3,1} + \tilde{\sigma}_b\mu_{3,0},\\ \dot{\mu}_{4,0} &= 4\rho_u B(\mu_{3,0} + 3B\mu_{1,0} + 6B^3\mu_{1,0} + 6B^3 g_0) - 4d\mu_{4,0} + \tilde{\sigma}_u\mu_{4,1} - \tilde{\sigma}_b\mu_{4,0},\\ \dot{\mu}_{4,1} &= 4\rho_b B(\mu_{3,1} + 3B\mu_{1,1} + 6B^3\mu_{1,1} + 6B^3 g_1) - 4d\mu_{4,1} - \tilde{\sigma}_u\mu_{4,1} + \tilde{\sigma}_b\mu_{4,0}.\end{aligned} \tag{9}$$

When $h = 2$ or $h = 3$, inserting Eq. (8) into Eqs. (6) and (9) yields a set of closed moment equations, from which we can obtain the values of $g_i$, $\mu_{1,i}$, $\mu_{2,i}$, $\mu_{3,i}$ and $\mu_{4,i}$. Finally we can use Eq. (7) to determine the values of $\tilde{\sigma}_b$ and $\tilde{\sigma}_u$.

#### Algorithm for the 4-HM

We next consider the 4-HM, which maps the nonlinear network to the following linear one:

$$G \xrightarrow{\bar{\sigma}_b} G^*, \quad G^* \xrightarrow{\bar{\sigma}_u} G,$$
$$G \xrightarrow{\bar{\rho}_u} G + kP, \quad G^* \xrightarrow{\bar{\rho}_b} G^* + kP, \quad P \xrightarrow{d} \varnothing,$$

where $\bar{\sigma}_b$ and $\bar{\sigma}_u$ are the effective rates of gene state switching, and $\bar{\rho}_u$ and $\bar{\rho}_b$ are the effective burst frequencies in the two gene states. Next we determine the values of the four effective parameters. For the linear network, similar computations show that the evolution of zeroth, first, and second-order moments are given by

$$\begin{aligned}
\dot{g}_0 &= \bar{\sigma}_u g_1 - \bar{\sigma}_b g_0, \\
\dot{\mu}_{1,0} &= \bar{\rho}_u B g_0 - d\mu_{1,0} + \bar{\sigma}_u \mu_{1,1} - \bar{\sigma}_b \mu_{1,0}, \\
\dot{\mu}_{1,1} &= \bar{\rho}_b B g_1 - d\mu_{1,1} - \bar{\sigma}_u \mu_{1,1} + \bar{\sigma}_b \mu_{1,0}, \\
\dot{\mu}_{2,0} &= 2\bar{\rho}_u B(\mu_{1,0} + B g_0) - 2d\mu_{2,0} + \bar{\sigma}_u \mu_{2,1} - \bar{\sigma}_b \mu_{2,0}, \\
\dot{\mu}_{2,1} &= 2\bar{\rho}_b B(\mu_{1,1} + B g_1) - 2d\mu_{2,1} - \bar{\sigma}_u \mu_{2,1} + \bar{\sigma}_b \mu_{2,0}.
\end{aligned} \tag{10}$$

The four effective parameters $\bar{\sigma}_u$, $\bar{\sigma}_b$, $\bar{\rho}_u$, and $\bar{\rho}_b$ are chosen so that the two systems have the same zero, first, and second-order moments. Matching the last four identities in Eqs. (4) and (10), we find that $\bar{\rho}_b$ and $\bar{\rho}_u$ should be determined by the following system of linear equations:

$$\begin{aligned}
&\bar{\rho}_u B g_0 + \bar{\rho}_b B g_1 = \rho_u B g_0 + \rho_b B g_1 + h\sigma_u g_1 - h\sigma_b \mu_{h,0}, \\
&\bar{\rho}_u B(\mu_{1,0} + B g_0) + \bar{\rho}_b B(\mu_{1,1} + B g_1) = \rho_u B(\mu_{1,0} + B g_0) + \rho_b B(\mu_{1,1} + B g_1) \\
&\qquad + h\sigma_u \mu_{1,1} + \frac{h(h-1)}{2}\sigma_u g_1 - h\sigma_b \mu_{h+1,0} - \frac{h(h-1)}{2}\sigma_b \mu_{h,0}.
\end{aligned} \tag{11}$$

Similarly, matching the first and third identities in Eqs. (4) and (10), we find that $\bar{\sigma}_b$ and $\bar{\sigma}_u$ should be determined by the following system of linear equations:

$$\begin{aligned}
&\bar{\sigma}_u g_1 - \bar{\sigma}_b g_0 = \sigma_u g_1 - \sigma_b \mu_{h,0}, \\
&\bar{\sigma}_u \mu_{1,1} - \bar{\sigma}_b \mu_{1,0} = \sigma_u \mu_{1,1} - \sigma_b \mu_{h+1,0} + (\bar{\rho}_b - \rho_b) B g_1,
\end{aligned} \tag{12}$$

where $\bar{\rho}_b$ has been determined using Eq. (11). In the case of non-cooperative binding ($h = 1$), inserting Eqs. (11) and (12) into Eq. (10) yields a set of closed moment equations, i.e.

$$\begin{aligned}
\dot{g}_0 &= \bar{\sigma}_u(g_i, \mu_{1,i}, \mu_{2,i}) g_1 - \bar{\sigma}_b(g_i, \mu_{1,i}, \mu_{2,i}) g_0, \\
\dot{\mu}_{1,0} &= \bar{\rho}_u(g_i, \mu_{1,i}, \mu_{2,i}) B g_0 - d\mu_{1,0} + \bar{\sigma}_u(g_i, \mu_{1,i}, \mu_{2,i}) \mu_{1,1} - \bar{\sigma}_b(g_i, \mu_{1,i}, \mu_{2,i}) \mu_{1,0}, \\
\dot{\mu}_{1,1} &= \bar{\rho}_b(g_i, \mu_{1,i}, \mu_{2,i}) B g_1 - d\mu_{1,1} - \bar{\sigma}_u(g_i, \mu_{1,i}, \mu_{2,i}) \mu_{1,1} + \bar{\sigma}_b(g_i, \mu_{1,i}, \mu_{2,i}) \mu_{1,0}, \\
\dot{\mu}_{2,0} &= 2\bar{\rho}_u(g_i, \mu_{1,i}, \mu_{2,i}) B(\mu_{1,0} + B g_0) - 2d\mu_{2,0} + \bar{\sigma}_u(g_i, \mu_{1,i}, \mu_{2,i}) \mu_{2,1} - \bar{\sigma}_b(g_i, \mu_{1,i}, \mu_{2,i}) \mu_{2,0}, \\
\dot{\mu}_{2,1} &= 2\bar{\rho}_b(g_i, \mu_{1,i}, \mu_{2,i}) B(\mu_{1,1} + B g_1) - 2d\mu_{2,1} - \bar{\sigma}_u(g_i, \mu_{1,i}, \mu_{2,i}) \mu_{2,1} + \bar{\sigma}_b(g_i, \mu_{1,i}, \mu_{2,i}) \mu_{2,0}.
\end{aligned}$$

from which one can obtain the approximate values of $g_i$, $\mu_{1,i}$, and $\mu_{2,i}$. Finally we can use Eqs. (11) and (12) to determine the values of $\bar{\rho}_u$, $\bar{\rho}_b$, $\bar{\sigma}_u$, and $\bar{\sigma}_b$.

Alternatively, when $h = 1$, combining the first three identities in Eq. (4) and the last two identities in Eq. (10), and then inserting Eqs. (11) and (12) into Eq. (10) also yield a set of closed moment equations, i.e.

$$\begin{aligned}
\dot{g}_0 &= \sigma_u g_1 - \sigma_b \mu_{1,0}, \\
\dot{\mu}_{1,0} &= \rho_u B g_0 - d\mu_{1,0} + \sigma_u(\mu_{1,1} + g_1) - \sigma_b(\mu_{2,0} + \mu_{1,0}), \\
\dot{\mu}_{1,1} &= \rho_b B g_1 - d\mu_{1,1} - \sigma_u \mu_{1,1} + \sigma_b \mu_{2,0}, \\
\dot{\mu}_{2,0} &= 2\bar{\rho}_u(g_i, \mu_{1,i}, \mu_{2,i}) B(\mu_{1,0} + B g_0) - 2d\mu_{2,0} + \bar{\sigma}_u(g_i, \mu_{1,i}, \mu_{2,i}) \mu_{2,1} - \tilde{\sigma}_b(g_i, \mu_{1,i}, \mu_{2,i}) \mu_{2,0}, \\
\dot{\mu}_{2,1} &= 2\bar{\rho}_b(g_i, \mu_{1,i}, \mu_{2,i}) B(\mu_{1,1} + B g_1) - 2d\mu_{2,1} - \bar{\sigma}_u(g_i, \mu_{1,i}, \mu_{2,i}) \mu_{2,1} + \tilde{\sigma}_b(g_i, \mu_{1,i}, \mu_{2,i}) \mu_{2,0}.
\end{aligned}$$

In the case of cooperative binding ($h \geq 2$), we need higher-order moment equations. For example, direct computations show that the evolution of third and fourth moments are given by

$$\begin{aligned}
\dot{\mu}_{3,0} &= 3\bar{\rho}_u B(\mu_{2,0} + 2B\mu_{1,0} + 2B^2 g_0) - 3d\mu_{3,0} + \bar{\sigma}_u\mu_{3,1} - \bar{\sigma}_b\mu_{3,0}, \\
\dot{\mu}_{3,1} &= 3\bar{\rho}_b B(\mu_{2,1} + 2B\mu_{1,1} + 2B^2 g_1) - 3d\mu_{3,1} - \bar{\sigma}_u\mu_{3,1} + \bar{\sigma}_b\mu_{3,0}, \\
\dot{\mu}_{4,0} &= 4\bar{\rho}_u B(\mu_{3,0} + 3B\mu_{1,0} + 6B^3\mu_{1,0} + 6B^3 g_0) - 4d\mu_{4,0} + \bar{\sigma}_u\mu_{4,1} - \bar{\sigma}_b\mu_{4,0}, \\
\dot{\mu}_{4,1} &= 4\bar{\rho}_b B(\mu_{3,1} + 3B\mu_{1,1} + 6B^3\mu_{1,1} + 6B^3 g_1) - 4d\mu_{4,1} - \bar{\sigma}_u\mu_{4,1} + \bar{\sigma}_b\mu_{4,0}.
\end{aligned} \tag{13}$$

When $h = 2$ or $h = 3$, inserting Eqs. (11) and (12) into Eqs. (10) and (13) yields a set of closed moment equations, from which we can obtain the values of $g_i$, $\mu_{1,i}$, $\mu_{2,i}$, $\mu_{3,i}$ and $\mu_{4,i}$. Finally we can use Eqs. (11) and (12) to determine the values of $\bar{\rho}_u$, $\bar{\rho}_b$, $\bar{\sigma}_u$, and $\bar{\sigma}_b$. As stated in the main text, the 4-HM may lead to numerical instability when computing the time-dependent distributions. This is the main limitation of our moment closure technique.

### Note S3. Number of moment equations to be solved for a general network

Consider a general regulatory network involving $M$ distinct genes, each of which can be in two states: an inactive state $G_j$ and an active state $G_j^*$. The protein associated with gene $G_j$ is denoted by $P_j$. For simplicity, we assume that each protein-gene binding reaction has the same cooperativity $h$. For Holimap, it is difficult to compute the exact number of moment equations $L$ to be solved since $L$ depends on the intricate regulatory reactions of the network. However, we can estimate the lower and upper bounds of $L$.

For an autoregulatory gene circuit, we have seen that $L = 3 + 2h$ for Holimap. For a general network, we need to solve at least $3 + 2h$ moment equations for each gene state. Since there are two states for each gene, the total number of gene states for the network is $2^M$. Hence the total number of moment equations to be solved has the following lower bound:

$$L \geq 2^M(3 + 2h). \tag{14}$$

On the other hand, given the cooperativity $h$, the Holimap algorithm for finding the effective parameters requires the solution of $(h + 1)$-order moment equations. For each gene state, it is easy to see that the number of $m$th-order mixed moment is $C_{M+m-1,m}$, where $C_{n,m} = n!/m!(n - m)!$ is the combinatorial number. This can be seen as follows. It is clear that each mixed $m$th-order moment can be represented by $\langle n_1^{x_1} n_2^{x_2} \cdots n_M^{x_M} \rangle$, where $n_j$ denotes the number of protein $P_j$ and $x_1, x_2, \cdots, x_M$ are integers satisfying

$$x_1 + x_2 + \cdots + x_M = m, \quad x_1, x_2, \cdots, x_M \geq 0.$$

It is easy to check that the number of $M$-tuples $(x_1, x_2, \cdots, x_M)$ satisfying the above equation is $C_{M+m-1,m}$ and hence the number of $m$th-order mixed moment is $C_{M+m-1,m}$ for each gene state. Hence for a general network, the total number of moment equations to be solved has the following upper bound:

$$L \leq 2^M \sum_{m=0}^{h+1} C_{M+m-1,m} = 2^M(C_{M-1,0} + C_{M,1} + \cdots + C_{M+h,h+1}). \tag{15}$$

Combining Eqs. (14) and (15), it can be seen that

$$2^M(3 + 2h) \leq L \leq 2^M \sum_{m=0}^{h+1} C_{M+m-1,m}. \tag{16}$$

Simple algebra shows that $\sum_{m=0}^{h+1} C_{M+m-1,m}$ is a polynomial of $M$ of degree $h$ and is also a polynomial of $h$ of degree $M$. Hence it follows from Eq. (16) that $L$ scales with $M$ exponentially (this is mainly because the number of gene states scales with $M$ exponentially) and $L$ scales with $h$ polynomially.

#### Note S4. Deterministic rate equations for 2-node and 3-node networks

For the 2-node network shown in Fig. 3(a) in the main text, the reaction scheme is given by

$$\begin{aligned}
G_1 + h_2P_2 &\xrightarrow{\sigma_{b1}} G_1^*, \quad G_1^* \xrightarrow{\sigma_{u1}} G_1 + h_2P_2,\\
G_2 + h_1P_1 &\xrightarrow{\sigma_{b2}} G_2^*, \quad G_2^* \xrightarrow{\sigma_{u2}} G_2 + h_1P_1,\\
G_1 &\xrightarrow{\rho_{u1}} G_1 + P_1, \quad G_1^* \xrightarrow{\rho_{b1}} G_1^* + P_1,\\
G_2 &\xrightarrow{\rho_{u2}} G_2 + P_2, \quad G_2^* \xrightarrow{\rho_{b2}} G_2^* + P_2,\\
P_1 &\xrightarrow{d_1} \varnothing, \quad P_2 \xrightarrow{d_2} \varnothing.
\end{aligned}$$

Here the genes $G_1$ and $G_2$ regulate each other. Clearly, $G_2$ positively regulates $G_1$ if $\rho_{b1} > \rho_{u1}$, while $G_2$ negatively regulates $G_1$ if $\rho_{b1} < \rho_{u1}$; similarly, $G_1$ positively regulates $G_2$ if $\rho_{b2} > \rho_{u2}$, while $G_1$ negatively regulates $G_2$ if $\rho_{b2} < \rho_{u2}$. Deterministically, based on the law of mass action, the network can be modelled by the following ODEs:

$$\begin{aligned}
\dot{w}_1 &= \sigma_{b1}x_2^{h_2}(1-w_1) - \sigma_{u1}w_1,\\
\dot{w}_2 &= \sigma_{b2}x_1^{h_1}(1-w_2) - \sigma_{u2}w_2,\\
\dot{x}_1 &= \rho_{u1}(1-w_1) + \rho_{b1}w_1 - d_1x_1 + h_1[\sigma_{u2}w_2 - \sigma_{b2}x_1^{h_1}(1-w_2)],\\
\dot{x}_2 &= \rho_{u2}(1-w_2) + \rho_{b2}w_2 - d_2x_2 + h_2[\sigma_{u1}w_1 - \sigma_{b1}x_2^{h_2}(1-w_1)],
\end{aligned} \tag{17}$$

where $w_i$ and $x_i$ are the mean copy numbers of gene $G_i$ and protein $P_i$, respectively. In the adiabatic (fast switching) limit, the gene switching dynamics will rapidly reach equilibrium, i.e. $\dot{w}_1 = \dot{w}_2 = 0$, which implies that

$$w_1 = \frac{\sigma_{b1}x_2^{h_2}}{\sigma_{u1} + \sigma_{b1}x_2^{h_2}}, \quad w_2 = \frac{\sigma_{b2}x_1^{h_1}}{\sigma_{u2} + \sigma_{b2}x_1^{h_1}}. \tag{18}$$

Inserting Eq. (18) into Eq. (17) yields the well-known form of the deterministic rate equations for the time-evolution of the mean protein numbers:

$$\begin{aligned}
\dot{x}_1 &= \frac{\rho_{u1}K_1 + \rho_{b1}x_2^{h_2}}{K_1 + x_2^{h_2}} - d_1x_1,\\
\dot{x}_2 &= \frac{\rho_{u2}K_2 + \rho_{b2}x_1^{h_1}}{K_2 + x_1^{h_1}} - d_2x_2,
\end{aligned}$$

where $K_i = \sigma_{ui}/\sigma_{bi}$.

Similarly, for the 3-node network shown in Fig. 4(a) in the main text, the reaction scheme is given by

$$\begin{aligned}
G_1 + h_3P_3 &\xrightarrow{\sigma_{b1}} G_1^*, \quad G_1^* \xrightarrow{\sigma_{u1}} G_1 + h_3P_3,\\
G_2 + h_1P_1 &\xrightarrow{\sigma_{b2}} G_2^*, \quad G_2^* \xrightarrow{\sigma_{u2}} G_2 + h_1P_1,\\
G_3 + h_2P_2 &\xrightarrow{\sigma_{b3}} G_3^*, \quad G_3^* \xrightarrow{\sigma_{u3}} G_3 + h_2P_2,\\
G_1 &\xrightarrow{\rho_{u1}} G_1 + P_1, \quad G_1^* \xrightarrow{\rho_{b1}} G_1^* + P_1,\\
G_2 &\xrightarrow{\rho_{u2}} G_2 + P_2, \quad G_2^* \xrightarrow{\rho_{b2}} G_2^* + P_2,\\
G_3 &\xrightarrow{\rho_{u3}} G_3 + P_3, \quad G_3^* \xrightarrow{\rho_{b3}} G_3^* + P_3,\\
P_1 &\xrightarrow{d_1} \varnothing, \quad P_2 \xrightarrow{d_2} \varnothing, \quad P_3 \xrightarrow{d_3} \varnothing.
\end{aligned}$$

Here three genes $G_1$, $G_2$, and $G_3$ regulate each other in a cyclic manner. Deterministically, based on the law of mass action, the network can be modelled by the following ODEs:

$$\begin{aligned}
\dot{w}_1 &= \sigma_{b1}x_3^{h_3}(1-w_1) - \sigma_{u1}w_1, \\
\dot{w}_2 &= \sigma_{b2}x_1^{h_1}(1-w_2) - \sigma_{u2}w_2, \\
\dot{w}_3 &= \sigma_{b3}x_2^{h_2}(1-w_3) - \sigma_{u3}w_3, \\
\dot{x}_1 &= \rho_{u1}(1-w_1) + \rho_{b1}w_1 - d_1x_1 + h_1[\sigma_{u2}w_2 - \sigma_{b2}x_1^{h_1}(1-w_2)], \\
\dot{x}_2 &= \rho_{u2}(1-w_2) + \rho_{b2}w_2 - d_2x_2 + h_2[\sigma_{u3}w_3 - \sigma_{b3}x_2^{h_2}(1-w_3)], \\
\dot{x}_3 &= \rho_{u3}(1-w_3) + \rho_{b3}w_3 - d_3x_3 + h_3[\sigma_{u1}w_1 - \sigma_{b1}x_3^{h_3}(1-w_1)],
\end{aligned} \tag{19}$$

where $w_i$ and $x_i$ is the mean copy numbers of gene $G_i$ and protein $P_i$, respectively. In the adiabatic (fast switching) limit, the gene switching dynamics will rapidly reach equilibrium, i.e. $\dot{w}_1 = \dot{w}_2 = \dot{w}_3 = 0$, which implies that

$$w_1 = \frac{\sigma_{b1}x_3^{h_3}}{\sigma_{u1} + \sigma_{b1}x_3^{h_3}}, \quad w_2 = \frac{\sigma_{b2}x_1^{h_1}}{\sigma_{u2} + \sigma_{b2}x_1^{h_1}}, \quad w_3 = \frac{\sigma_{b3}x_2^{h_2}}{\sigma_{u3} + \sigma_{b3}x_2^{h_2}}. \tag{20}$$

Inserting Eq. (20) into Eq. (19) yields the well-known form of the deterministic rate equations for the time-evolution of the mean protein numbers:

$$\begin{aligned}
\dot{x}_1 &= \frac{\rho_{u1}K_1 + \rho_{b1}x_3^{h_3}}{K_1 + x_3^{h_3}} - d_1x_1, \\
\dot{x}_2 &= \frac{\rho_{u2}K_2 + \rho_{b2}x_1^{h_1}}{K_2 + x_1^{h_1}} - d_2x_2, \\
\dot{x}_2 &= \frac{\rho_{u3}K_3 + \rho_{b3}x_2^{h_2}}{K_3 + x_2^{h_2}} - d_3x_3,
\end{aligned}$$

where $K_i = \sigma_{ui}/\sigma_{bi}$.

### Note S5. Applications of Holimap to networks with multi-transcription factor interactions

We consider a two-node regulatory network where gene $G_2$ is regulated by gene $G_1$, and gene $G_1$ is regulated by both gene $G_2$ and itself (Supplementary Fig. 6(a)). Specifically, we assume that the promoter of gene $G_1$ has two binding sites: one for protein $P_1$ and the other for protein $P_2$. According to whether these two binding sites are occupied or not, there are four conformal states for gene $G_1$ with four binding rates $\lambda_{b1}$, $\lambda'_{b1}$, $\sigma_{b1}$, and $\sigma'_{b1}$, as well as four unbinding rates $\lambda_{u1}$, $\lambda'_{u1}$, $\sigma_{u1}$, and $\sigma'_{u1}$. These four gene states are denoted by $G_1^{uu}$, $G_1^{bu}$, $G_1^{ub}$, and $G_1^{bb}$. Here $G_1^{bu}$ represents the state where protein $P_1$ is bound and protein $P_2$ is unbound; the other states can be understood similarly. The synthesis rates for the four gene states are denoted by $\rho_{uu1}$, $\rho_{bu1}$, $\rho_{ub1}$, and $\rho_{bb1}$.

To further simplify the model, we make two additional assumptions. First, we assume that the binding and unbinding rates satisfy $\lambda_{b1}\sigma'_{b1}\lambda'_{u1}\sigma_{u1} = \sigma_{b1}\lambda'_{b1}\sigma'_{u1}\lambda_{u1}$; this is called the Wegscheider condition, which guarantees that gene switching dynamics is in detailed balance [6]. Second, we assume that the synthesis rates for the four gene states can only take two different values — only one state is endowed with a high synthesis rate $\rho_H$ and the synthesis rates $\rho_L$ for the other states are lower (this is an example of the AND-logic gate [7]). This ensures that the protein distribution has at most two modes. For example, if gene $G_1$ is activated by both gene $G_2$ and itself, then the gene state $G_1^{bb}$ has the highest synthesis rate since the two binding sites are both occupied. In this case, we have

$$\rho_{bb} = \rho_H, \quad \rho_{uu} = \rho_{bu} = \rho_{ub} = \rho_L.$$

If the dynamics of gene switching is much faster than the dynamics of protein synthesis and degradation (adiabatic limit), then the evolution of the abundances of the two proteins is governed by the following deterministic system:

$$\dot{x}_1 = \frac{\rho_{uu1} + \rho_{bu1}x_1/K + \rho_{ub1}x_2/L + \rho_{bb1}x_1x_2/M}{1 + x_1/K + x_2/L + x_1x_2/M} - d_1x_1,$$
$$\dot{x}_2 = \frac{\rho_{u2}\sigma_{u2} + \rho_{b2}\sigma_{b2}x_1}{\sigma_{u2} + \sigma_{b2}x_1} - d_2x_2,$$

where $K$, $L$, and $M$ are constants defined by

$$K = \frac{\lambda_{u1}}{\lambda_{b1}}, \quad L = \frac{\sigma_{u1}}{\sigma_{b1}}, \quad M = K\frac{\sigma'_{u1}}{\sigma'_{b1}} = L\frac{\lambda'_{u1}}{\lambda'_{b1}}. \tag{21}$$

Note that the above deterministic rate equations correspond to those proposed in Ref. [6].

For this network, we can also use Holimap to transform it into a four-state linear network (Supplementary Fig. 6(b)), whose protein distribution can be solved analytically both in steady state and in time [8, 9]. We devise two types of Holimaps. The first type is the four-parameter Holimap (4-HM), which replaces the four binding rates by four effective parameters. Since there are four binding rates for gene $G_1$ in the current model, the 4-HM considered here should be viewed as the counterpart of the LMA in previous models, which is the conventional approach. The second type of Holimap modifies the four binding rates and four unbinding rates simultaneously; this yields an eight-parameter Holimap (8-HM). Simulations show that the 4-HM typically accurately predicts unimodal protein distributions, while the 8-HM outperforms it when the distributions are bimodal (Supplementary Fig. 6(c)).

Note that the network shown in Supplementary Fig. 6(a) can describe 12 distinct topologies (Supplementary Fig. 6(d)), according to the sign of feedback controls. These 12 topologies can be further classified into two groups according to whether the inputs of gene $G_1$ are coherent or incoherent. Here coherent (incoherent) inputs mean that the autoregulation of gene $G_1$ and its regulation by gene $G_2$ have the same (opposite) signs. Moreover, for each topology, the binding sites for the two proteins in the promoter region of gene $G_1$ can be of three types: independent binding ($M = KL$), positive cooperative binding ($M < KL$), and negative cooperative binding ($M > KL$) [10], where $K$, $L$, and $M$ are constants defined in Eq. (21). Here independent binding means that the binding of one protein does not affect the binding of the other protein. Positive (negative) cooperativity means that protein binding at one site increases (decreases) protein binding at the other site.

We next study how various network properties influence the shape of the protein distribution. For each topology and each type of interaction, we randomly selected $10^4$ sets of model parameters, and computed the number of parameter sets $Q$ that result in a bimodal steady-state protein distribution. The value of $Q$ reflects the possibility that the system displays bimodality and it is called the $Q$-value in previous studies [11]. Interestingly, we find that for all the 6 networks with coherent (incoherent) inputs, positive (negative) cooperativity in the promoter region gives rise to a higher $Q$-value (Supplementary Fig. 6(d)). To further test Holimap, we compare the real $Q$-value computed using FSP (shown by solid bars) with those predicted by the 4-HM and 8-HM (shown by red and blue lines, respectively). It is clear that independent of the type of network topology, the type of inputs, and the type of binding, the 8-HM excellently predicts the $Q$-value, while the conventional 4-HM significantly underestimates this value since it does not accurately capture the distribution shape in the region where bimodality is manifest.

### Note S6. Estimation of the computational time for FSP

Because FSP depends on matrix exponentiation, its computational time is $O(N^3)$ with $N$ being the total truncation size, i.e. the total number of states in the truncated state space. For a three-node network, there are a total of $2^3 = 8$ gene states and hence the total truncation size is given by $N = 8N_1N_2N_3$, where $N_i$ is the truncation size for protein $P_i$. If we choose $N_i = 10$, then the running time for FSP is about 1.25 min, according to simulations. However, given the range of the distributions in Fig. 4(d) in the main text, it is clear that $N_i$ should be chosen to be at least 120. Hence in order to obtain an accurate protein distribution, the running time for FSP should be at least $1.25 \times (12 \times 12 \times 12)^3$ min, which is about $1.23 \times 10^4$ years.

### Note S7. Effective parameters for post-transcriptional and post-translational networks

#### Protein sequestration

We now consider the following gene regulatory network (Fig. 6(a) in the main text):

$$
\begin{gathered}
G_1 + h_1P_1 \xrightarrow{\sigma_{b1}} G_1^*, \quad G_1^* \xrightarrow{\sigma_{u1}} G_1 + h_1P_1, \\
G_2 + h_2P_2 \xrightarrow{\sigma_{b2}} G_2^*, \quad G_2^* \xrightarrow{\sigma_{u2}} G_2 + h_2P_2, \\
G_1 \xrightarrow{\rho_{u1}} G_1 + P_1, \quad G_1^* \xrightarrow{\rho_{b1}} G_1^* + P_1, \\
G_2 \xrightarrow{\rho_{u2}} G_2 + P_2, \quad G_2^* \xrightarrow{\rho_{b2}} G_2^* + P_2, \\
P_1 \xrightarrow{d_1} \varnothing, \quad P_2 \xrightarrow{d_2} \varnothing, \quad P_1 + P_2 \xrightarrow{\alpha} C.
\end{gathered}
$$

Here protein $P_i$ expressed from gene $G_i$ regulates its own transcription. Moreover, proteins $P_1$ and $P_2$ can bind to each other and form an inactive complex $C$. The microstate of the system can be represented by the ordered four-tuple $(i_1, i_2, n_1, n_2)$, where $n_k$ is the number of protein $P_k$ and $i_k$ is the state of gene $G_k$ with $i_k = 0, 1$ corresponding to the unbound and bound states, respectively. Let $p_{i_1,i_2,n_1,n_2}$ denote the probability of the system being in microstate $(i_1, i_2, n_1, n_2)$. Then the dynamics of the system is governed by the CMEs

$$
\begin{aligned}
\dot{p}_{0,0,n_1,n_2} = {} & \rho_{u1}[p_{0,0,n_1-1,n_2} - p_{0,0,n_1,n_2}] + d_1[(n_1+1)p_{0,0,n_1+1,n_2} - n_1 p_{0,0,n_1,n_2}] \\
& + \rho_{u2}[p_{0,0,n_1,n_2-1} - p_{0,0,n_1,n_2}] + d_2[(n_2+1)p_{0,0,n_1,n_2+1} - n_2 p_{0,0,n_1,n_2}] \\
& + \alpha[(n_1+1)(n_2+1)p_{0,0,n_1+1,n_2+1} - n_1 n_2 p_{0,0,n_1,n_2}] \\
& + \left[\sigma_{u1} p_{1,0,n_1-h_1,n_2} - \sigma_{b1}\frac{n_1!}{(n_1-h_1)!} p_{0,0,n_1,n_2}\right] \\
& + \left[\sigma_{u2} p_{0,1,n_1,n_2-h_2} - \sigma_{b2}\frac{n_2!}{(n_2-h_2)!} p_{0,0,n_1,n_2}\right], \\
\dot{p}_{0,1,n_1,n_2} = {} & \rho_{u1}[p_{0,1,n_1-1,n_2} - p_{0,1,n_1,n_2}] + d_1[(n_1+1)p_{0,1,n_1+1,n_2} - n_1 p_{0,1,n_1,n_2}] \\
& + \rho_{b2}[p_{0,1,n_1,n_2-1} - p_{0,1,n_1,n_2}] + d_2[(n_2+1)p_{0,1,n_1,n_2+1} - n_2 p_{0,1,n_1,n_2}] \\
& + \alpha[(n_1+1)(n_2+1)p_{0,1,n_1+1,n_2+1} - n_1 n_2 p_{0,1,n_1,n_2}] \\
& + \left[\sigma_{u1} p_{1,1,n_1-h_1,n_2} - \sigma_{b1}\frac{n_1!}{(n_1-h_1)!} p_{0,1,n_1,n_2}\right] \\
& + \left[\sigma_{b2}\frac{(n_2+h_2)!}{n_2!} p_{0,0,n_1,n_2+h_2} - \sigma_{u2} p_{0,1,n_1,n_2}\right], \\
\dot{p}_{1,0,n_1,n_2} = {} & \rho_{b1}[p_{1,0,n_1-1,n_2} - p_{1,0,n_1,n_2}] + d_1[(n_1+1)p_{1,0,n_1+1,n_2} - n_1 p_{1,0,n_1,n_2}] \\
& + \rho_{u2}[p_{1,0,n_1,n_2-1} - p_{1,0,n_1,n_2}] + d_2[(n_2+1)p_{1,0,n_1,n_2+1} - n_2 p_{1,0,n_1,n_2}] \\
& + \alpha[(n_1+1)(n_2+1)p_{1,0,n_1+1,n_2+1} - n_1 n_2 p_{1,0,n_1,n_2}] \\
& + \left[\sigma_{b1}\frac{(n_1+h_1)!}{n_1!} p_{0,0,n_1+h_1,n_2} - \sigma_{u1} p_{1,0,n_1,n_2}\right] \\
& + \left[\sigma_{u2} p_{1,1,n_1,n_2-h_2} - \sigma_{b2}\frac{n_2!}{(n_2-h_2)!} p_{1,0,n_1,n_2}\right], \\
\dot{p}_{1,1,n_1,n_2} = {} & \rho_{b1}[p_{1,1,n_1-1,n_2} - p_{1,1,n_1,n_2}] + d_1[(n_1+1)p_{1,1,n_1+1,n_2} - n_1 p_{1,1,n_1,n_2}] \\
& + \rho_{b2}[p_{1,1,n_1,n_2-1} - p_{1,1,n_1,n_2}] + d_2[(n_2+1)p_{1,1,n_1,n_2+1} - n_2 p_{1,1,n_1,n_2}] \\
& + \alpha[(n_1+1)(n_2+1)p_{1,1,n_1+1,n_2+1} - n_1 n_2 p_{1,1,n_1,n_2}] \\
& + \left[\sigma_{b1}\frac{(n_1+h_1)!}{n_1!} p_{0,1,n_1+h_1,n_2} - \sigma_{u1} p_{1,1,n_1,n_2}\right] \\
& + \left[\sigma_{b2}\frac{(n_2+h_2)!}{n_2!} p_{1,0,n_1,n_2+h_2} - \sigma_{u2} p_{1,1,n_1,n_2}\right].
\end{aligned}
$$

We next define the generating function

$$F_{i_1,i_2}(x,y) = \sum_{n_1,n_2=0}^{\infty} p_{i_1,i_2,n_1,n_2} x^{n_1} y^{n_2}.$$

Then the CMEs can be converted into the PDEs

$$\begin{aligned}
\partial_t F_{0,0} &= \rho_{u1}(x-1)F_{0,0} - d_1(x-1)\partial_x F_{0,0} + \rho_{u2}(y-1)F_{0,0} - d_2(y-1)\partial_y F_{0,0} - \alpha(xy-1)\partial_{xy}F_{0,0} \\
&\quad + \sigma_{u1}x^{h_1}F_{1,0} - \sigma_{b1}x^{h_1}\partial_x^{(h_1)}F_{0,0} + \sigma_{u2}y^{h_2}F_{0,1} - \sigma_{b2}y^{h_2}\partial_y^{(h_2)}F_{0,0}, \\
\partial_t F_{0,1} &= \rho_{u1}(x-1)F_{0,1} - d_1(x-1)\partial_x F_{0,1} + \rho_{b2}(y-1)F_{0,1} - d_2(y-1)\partial_y F_{0,1} - \alpha(xy-1)\partial_{xy}F_{0,1} \\
&\quad + \sigma_{u1}x^{h_1}F_{1,1} - \sigma_{b1}x^{h_1}\partial_x^{(h_1)}F_{0,1} + \sigma_{b2}\partial_y^{(h_2)}F_{0,0} - \sigma_{u2}F_{0,1}, \\
\partial_t F_{1,0} &= \rho_{b1}(x-1)F_{1,0} - d_1(x-1)\partial_x F_{1,0} + \rho_{u2}(y-1)F_{1,0} - d_2(y-1)\partial_y F_{1,0} - \alpha(xy-1)\partial_{xy}F_{1,0} \\
&\quad + \sigma_{b1}\partial_x^{(h_1)}F_{0,0} - \sigma_{u1}F_{1,0} + \sigma_{u2}y^{h_2}F_{1,1} - \sigma_{b2}y^{h_2}\partial_y^{(h_2)}F_{1,0}, \\
\partial_t F_{1,1} &= \rho_{b1}(x-1)F_{1,1} - d_1(x-1)\partial_x F_{1,1} + \rho_{b2}(y-1)F_{1,1} - d_2(y-1)\partial_y F_{1,1} - \alpha(xy-1)\partial_{xy}F_{1,1} \\
&\quad + \sigma_{b1}\partial_x^{(h_1)}F_{0,1} - \sigma_{u1}F_{1,1} + \sigma_{b2}\partial_y^{(h_2)}F_{1,0} - \sigma_{u2}F_{1,1}.
\end{aligned}$$

We next only focus on the dynamics of protein $P_1$. Let

$$g_{i_1} = \sum_{i_2=0}^{1}\sum_{n_1,n_2=0}^{\infty} p_{i_1,i_2,n_1,n_2} = \sum_{i_2=0}^{1} F_{i_1,i_2}(1,1)$$

be the probability of gene $G_1$ being in state $i_1$ and let

$$\mu_{k,l,i_1} = \sum_{i_2=0}^{1}\sum_{n_1,n_2=0}^{\infty} \frac{n_1!n_2!}{(n_1-k)!(n_2-l)!} p_{i_1,i_2,n_1,n_2} = \sum_{i_2=0}^{1} \partial_x^{(k)}\partial_y^{(l)} F_{i_1,i_2}(1,1)$$

be the mixed factorial moment of protein numbers when gene $G_1$ is in state $i_1$. Straightforward computations show that the evolution of the zero and first moments are given by

$$\begin{aligned}
\dot{g}_0 &= \sigma_{u1}g_1 - \sigma_{b1}\mu_{h_1,0,0}, \\
\dot{\mu}_{1,0,0} &= \rho_{u1}g_0 - d_1\mu_{1,0,0} - \alpha\mu_{1,1,0} + \sigma_{u1}(\mu_{1,0,1} + h_1 g_1) - \sigma_{b1}(\mu_{h_1+1,0,0} + h_1\mu_{h_1,0,0}), \\
\dot{\mu}_{1,0,1} &= \rho_{b1}g_1 - d_1\mu_{1,0,1} - \alpha\mu_{1,1,1} + \sigma_{b1}\mu_{h_1+1,0,0} - \sigma_{u1}\mu_{1,0,1}.
\end{aligned} \tag{22}$$

We next use Holimap to map the nonlinear network to the following linear one:

$$G_1 \xrightarrow{\tilde{\sigma}_{b1}} G_1^*, \quad G_1^* \xrightarrow{\tilde{\sigma}_{u1}} G_1,$$

$$G_1 \xrightarrow{\rho_{u1}} G_1 + P_1, \quad G^* \xrightarrow{\rho_{b1}} G_1^* + P_1, \quad P_1 \xrightarrow{\tilde{d}_1} \varnothing.$$

The evolution of the zero and first moments for the linear network is given by

$$\begin{aligned}
\dot{g}_0 &= \tilde{\sigma}_{u1}g_1 - \tilde{\sigma}_{b1}g_0, \\
\dot{\mu}_{1,0,0} &= \rho_{u1}g_0 - \tilde{d}_1\mu_{1,0,0} + \tilde{\sigma}_{u1}\mu_{1,0,1} - \tilde{\sigma}_{b1}\mu_{1,0,0}, \\
\dot{\mu}_{1,0,1} &= \rho_{b1}g_1 - \tilde{d}_1\mu_{1,0,1} + \tilde{\sigma}_{b1}\mu_{1,0,0} - \tilde{\sigma}_{u1}\mu_{1,0,1}.
\end{aligned} \tag{23}$$

The three effective parameters $\tilde{\sigma}_{u1}$, $\tilde{\sigma}_{b1}$, and $\tilde{d}_1$ should be chosen so that the two systems have the same zero and first moments. Matching Eqs. (22) and (23), we find that $\tilde{d}$ should be determined by the following system of linear equations:

$$\tilde{d}_1\mu_{1,0,1} = d_1\mu_{1,0,1} + \alpha\mu_{1,1,1}. \tag{24}$$

Similarly, $\tilde{\sigma}_b$ and $\tilde{\sigma}_u$ should be determined by the following system of linear equations:

$$\begin{aligned}
\tilde{\sigma}_{u1}g_1 - \tilde{\sigma}_{b1}g_0 &= \sigma_{u1}g_1 - \sigma_{b1}\mu_{h_1,0,0}, \\
\tilde{\sigma}_{u1}\mu_{1,0,1} - \tilde{\sigma}_{b1}\mu_{1,0,0} &= \sigma_{u1}\mu_{1,0,1} - \sigma_{b1}\mu_{h_1+1,0,0}.
\end{aligned} \tag{25}$$

In summary, once we have computed the values of moments, we can use Eqs. (24) and (25) to determine the values of the three effective parameters.

#### Protein phosphorylation

We now consider the following gene regulatory network (Fig. 6(b) in the main text):

$$
\begin{gathered}
G + hP_2 \xrightarrow{\sigma_b} G^*, \quad G^* \xrightarrow{\sigma_u} G + hP_2, \\
G \xrightarrow{\rho_u} G + P, \quad G^* \xrightarrow{\rho_b} G^* + P, \\
P + E_1 \xrightarrow{a_1} C_1, \quad C_1 \xrightarrow{b_1} E_1 + P, \quad C_1 \xrightarrow{c_1} E_1 + P_1, \\
P_1 + E_2 \xrightarrow{a_2} C_2, \quad C_2 \xrightarrow{b_2} E_2 + P_1, \quad C_2 \xrightarrow{c_2} E_2 + P, \\
P_1 + E_3 \xrightarrow{a_3} C_3, \quad C_3 \xrightarrow{b_3} E_3 + P_1, \quad C_3 \xrightarrow{c_3} E_3 + P_2, \\
P_2 + E_4 \xrightarrow{a_4} C_4, \quad C_4 \xrightarrow{b_4} E_4 + P_2, \quad C_4 \xrightarrow{c_4} E_4 + P_1, \\
P \xrightarrow{d_0} \varnothing, \quad P_1 \xrightarrow{d_1} \varnothing, \quad P_2 \xrightarrow{d_2} \varnothing.
\end{gathered}
$$

Here the protein can be reversibly phosphorylated from the form $P$ into the forms $P_1$ and $P_2$, successively. The phosphorylated form $P_2$ of protein can bind to the gene and regulates its expression. Both phosphorylation and dephosphorylation are enzyme-catalyzed and are described using Michaelis-Menten kinetics, where $E_i$ are free enzymes and $C_i$ are enzyme-protein complexes. The total number of $E_i$ and $C_i$ is denoted by $N_i$, which is a constant.

The microstate of the system can be represented by the ordered eight-tuple $(i, n_0, n_1, n_2, m_1, m_2, m_3, m_4)$, where $n_0$ is the number of unphosphorylated protein $P$, $n_1$ and $n_2$ are the numbers of phosphorylated proteins $P_1$ and $P_2$, respectively, $m_k$ is the number of complex $C_k$, and $i$ is the state of the gene with $i = 0, 1$ corresponding to the unbound and bound states, respectively. Let $p_{i,n_0,n_1,n_2,m_1,m_2,m_3,m_4}$ denote the probability of the system being in microstate $(i, n_0, n_1, n_2, m_1, m_2, m_3, m_4)$. To proceed, let

$$
g_i = \sum_{n_0,n_1,n_2,m_1,m_2,m_3,m_4=0}^{\infty} p_{i,n_0,n_1,n_2,m_1,m_2,m_3,m_4}
$$

be the probability of the gene being in state $i$ and let

$$
\mu_{k_0,k_1,k_2,l_1,l_2,l_3,l_4,i} = \sum_{n_0,n_1,n_2,m_1,m_2,m_3,m_4=0}^{\infty} \prod_{\alpha=0}^{2} \frac{n_\alpha}{(n_\alpha - k_\alpha)!} \prod_{\beta=1}^{4} \frac{m_\beta}{(m_\beta - l_\beta)!} p_{i,n_0,n_1,n_2,m_1,m_2,m_3,m_4}
$$

be the mixed factorial moment of protein and complex numbers when the gene is in state $i$. Straightforward computations show that the evolution of the zero and first moments are given by

$$
\begin{aligned}
\dot{g}_0 &= \sigma_u g_1 - \sigma_b \mu_{0,0,h,0,0,0,0,0}, \\
\dot{g}_1 &= \sigma_b \mu_{0,0,h,0,0,0,0,0} - \sigma_u g_1, \\
\dot{\mu}_{1,0,0,0,0,0,0,0} &= \rho_u g_0 - (d_0 + a_1 N_1) \mu_{1,0,0,0,0,0,0,0} + a_1 \mu_{1,0,0,1,0,0,0,0} \\
&\quad + b_1 \mu_{0,0,0,1,0,0,0,0} + c_2 \mu_{0,0,0,0,1,0,0,0} + \sigma_u \mu_{1,0,0,0,0,0,0,1} - \sigma_b \mu_{1,0,h,0,0,0,0,0}, \\
\dot{\mu}_{1,0,0,0,0,0,0,1} &= \rho_b g_1 - (d_0 + a_1 N_1) \mu_{1,0,0,0,0,0,0,1} + a_1 \mu_{1,0,0,1,0,0,0,1} \\
&\quad + b_1 \mu_{0,0,0,1,0,0,0,1} + c_2 \mu_{0,0,0,0,1,0,0,1} + \sigma_b \mu_{1,0,h,0,0,0,0,0} - \sigma_u \mu_{1,0,0,0,0,0,0,1}.
\end{aligned} \tag{26}
$$

We next use Holimap to map the nonlinear network to the following linear one:

$$
\begin{gathered}
G \xrightarrow{\tilde{\sigma}_b} G^*, \quad G^* \xrightarrow{\tilde{\sigma}_u} G, \\
G \xrightarrow{\rho_u} G + P_0, \quad G^* \xrightarrow{\rho_b} G^* + P_0, \quad P_0 \xrightarrow{\tilde{d}_0} \varnothing.
\end{gathered}
$$

The evolution of the zero and first moments for the linear network is given by

$$
\begin{aligned}
\dot{g}_0 &= \tilde{\sigma}_u g_1 - \tilde{\sigma}_b g_0, \\
\dot{\mu}_{1,0,0,0,0,0,0,0} &= \rho_u g_0 - \tilde{d}_0 \mu_{1,0,0,0,0,0,0,0} + \tilde{\sigma}_u \mu_{1,0,0,0,0,0,0,1} - \tilde{\sigma}_b \mu_{1,0,0,0,0,0,0,0}, \\
\dot{\mu}_{1,0,0,0,0,0,0,1} &= \rho_b g_1 - \tilde{d}_0 \mu_{1,0,0,0,0,0,0,1} + \tilde{\sigma}_b \mu_{1,0,0,0,0,0,0,0} - \tilde{\sigma}_u \mu_{1,0,0,0,0,0,0,1}.
\end{aligned} \tag{27}
$$

The three effective parameters $\bar{\sigma}_u$, $\bar{\sigma}_b$, and $\bar{d}_0$ should be chosen so that the two systems have the same zero and first moments. Matching Eqs. (26) and (27), we find that $\bar{d}_0$ should be determined by the following system of linear equations:

$$\begin{aligned}\tilde{d}_0(\mu_{1,0,0,0,0,0,0,0,0} + \mu_{1,0,0,0,0,0,0,0,1}) = (d_0 + a_1N_1)(\mu_{1,0,0,0,0,0,0,0,0} + \mu_{1,0,0,0,0,0,0,0,1})\\ - a_1(\mu_{1,0,0,1,0,0,0,0} + \mu_{1,0,0,1,0,0,0,1})\\ - b_1(\mu_{0,0,0,1,0,0,0,0} + \mu_{0,0,0,1,0,0,0,1})\\ - c_2(\mu_{0,0,0,0,1,0,0,0} + \mu_{0,0,0,0,1,0,0,1}).\end{aligned} \tag{28}$$

This can be rewritten as

$$\tilde{d}_0\langle n_0\rangle = (d_0 + a_1N_1)\langle n_0\rangle - a_1\langle n_0m_1\rangle - b_1\langle m_1\rangle - c_2\langle m_2\rangle.$$

Similarly, $\bar{\sigma}_b$ and $\bar{\sigma}_u$ should be determined by the following system of linear equations:

$$\begin{aligned}&\tilde{\sigma}_u g_1 - \tilde{\sigma}_b g_0 = \sigma_u g_1 - \sigma_b\mu_{0,0,h,0,0,0,0,0},\\ &\tilde{\sigma}_u\mu_{1,0,0,0,0,0,0,1} - \tilde{\sigma}_b\mu_{1,0,0,0,0,0,0,0,0} = \sigma_u\mu_{1,0,0,0,0,0,0,1} - \sigma_b\mu_{1,0,h,0,0,0,0,0}.\end{aligned} \tag{29}$$

In summary, once we have computed the values of moments, we can use Eqs. (28) and (29) to determine the values of the three effective parameters.

#### Enzyme-catalyzed mRNA degradation

We now consider the following gene regulatory network (Fig. 6(d) in the main text):

$$\begin{gathered}G + hP \xrightarrow{\sigma_b} G^*,\quad G^* \xrightarrow{\sigma_u} G + hP,\\ G \xrightarrow{\rho_u} G + M,\quad G^* \xrightarrow{\rho_b} G^* + M,\\ E \xrightarrow{a} E^*,\quad E^* \xrightarrow{b} E,\\ M \xrightarrow{u} M + P,\quad P \xrightarrow{v} \varnothing,\\ M \xrightarrow{d} \varnothing,\quad E^* + M \xrightarrow{\alpha} E^*.\end{gathered}$$

Here the enzyme can convert between an active form $E^*$ and an inactive form $E$. The degradation of mRNA can occur spontaneously with rate $d$ and can be catalyzed by the active form $E^*$ of the enzyme with rate $\alpha$. The total number of $E$ and $E^*$ is denoted by $N$, which is a constant.

The microstate of the system can be represented by the ordered four-tuple $(i, m, n, l)$, where $m$ is the number of transcripts, $n$ is the number of protein, $l$ is the number of $E^*$, and $i$ is the state of the gene with $i = 0, 1$ corresponding to the unbound and bound states, respectively. Let $p_{i,m,n,l}$ denote the probability of the system being in microstate $(i, m, n, l)$. To proceed, let

$$g_i = \sum_{m,n,l=0}^{\infty} p_{i,m,n,l}$$

be the probability of the gene being in state $i$ and let

$$\mu_{k_m,k_n,k_l,i} = \sum_{m,n,l=0}^{\infty} \frac{m!n!l!}{(m-k_m)!(n-k_n)!(l-k_l)!}p_{i,m,n,l}$$

be the mixed factorial moment of mRNA, protein, and enzyme numbers when the gene is in state $i$. Straightforward computations show that the evolution of the zero and first moments are given by

$$\begin{aligned}&\dot{g}_0 = \sigma_u g_1 - \sigma_b\mu_{0,h,0,0},\\ &\dot{g}_1 = \sigma_b\mu_{0,h,0,0} - \sigma_u g_1,\\ &\dot{\mu}_{1,0,0,0} = \rho_u g_0 - d\mu_{1,0,0,0} - \alpha\mu_{1,0,1,0} + \sigma_u\mu_{1,0,0,1} - \sigma_b\mu_{1,h,0,0},\\ &\dot{\mu}_{1,0,0,1} = \rho_u g_1 - d\mu_{1,0,0,1} - \alpha\mu_{1,0,1,1} + \sigma_b\mu_{1,h,0,0} - \sigma_u\mu_{1,0,0,1}.\end{aligned} \tag{30}$$

We next use Holimap to map the nonlinear network to the following linear one:

$$G \xrightarrow{\tilde{\sigma}_b} G^*, \quad G^* \xrightarrow{\tilde{\sigma}_u} G,$$
$$G \xrightarrow{\rho_u} G + M, \quad G^* \xrightarrow{\rho_b} G^* + M, \quad M \xrightarrow{\tilde{d}} \varnothing.$$

The evolution of the zero and first moments for the linear network are given by

$$\begin{aligned} \dot{g}_0 &= \tilde{\sigma}_u g_1 - \tilde{\sigma}_b g_0, \\ \dot{\mu}_{1,0,0,0} &= \rho_u g_0 - \tilde{d}\mu_{1,0,0,0} + \bar{\sigma}_u \mu_{1,0,0,1} - \bar{\sigma}_b \mu_{1,0,0,0}, \\ \dot{\mu}_{1,0,0,1} &= \rho_b g_1 - \tilde{d}\mu_{1,0,0,1} + \bar{\sigma}_b \mu_{1,0,0,0} - \bar{\sigma}_u \mu_{1,0,0,1}. \end{aligned} \tag{31}$$

The three effective parameters $\tilde{\sigma}_u$, $\tilde{\sigma}_b$, and $\tilde{d}$ should be chosen so that the two systems have the same zero and first moments. Matching Eqs. (30) and (31), we find that $\tilde{d}$ should be determined by the following system of linear equations:

$$\tilde{d}(\mu_{1,0,0,0} + \mu_{1,0,0,1}) = d(\mu_{1,0,0,0} + \mu_{1,0,0,1}) + \alpha(\mu_{1,0,1,0} + \mu_{1,0,1,1}). \tag{32}$$

This can be rewritten as

$$\tilde{d}\langle m\rangle = d\langle m\rangle + \alpha\langle ml\rangle.$$

Similarly, $\bar{\sigma}_b$ and $\bar{\sigma}_u$ should be determined by the following system of linear equations:

$$\begin{aligned} \tilde{\sigma}_u g_1 - \tilde{\sigma}_b g_0 &= \sigma_u g_1 - \sigma_b \mu_{0,h,0,0}, \\ \tilde{\sigma}_u \mu_{1,0,0,1} - \tilde{\sigma}_b \mu_{1,0,0,0} &= \sigma_u \mu_{1,0,0,1} - \sigma_b \mu_{1,h,0,0}. \end{aligned} \tag{33}$$

In summary, once we have computed the values of moments, we can use Eqs. (32) and (33) to determine the values of the three effective parameters.

#### microRNA-mRNA interactions

We now consider the following gene regulatory network (Fig. 6(e) in the main text):

$$\begin{gathered} G_1 \xrightarrow{\sigma_{b1}} G_1^*, \quad G_1^* \xrightarrow{\sigma_{u1}} G_1, \\ G_2 \xrightarrow{\sigma_{b2}} G_2^*, \quad G_2^* \xrightarrow{\sigma_{u2}} G_2, \\ G_1 \xrightarrow{\rho_{u1}} G_1 + M, \quad G_1^* \xrightarrow{\rho_{b1}} G_1^* + M, \\ G_2 \xrightarrow{\rho_{u2}} G_2 + R, \quad G_2^* \xrightarrow{\rho_{b2}} G_2^* + R, \\ M + R \xrightarrow{\alpha} C_1, \quad C_1 \xrightarrow{\beta} M + R, \\ C_1 + R \xrightarrow{\alpha} C_2, \quad C_2 \xrightarrow{\beta} C_1 + R, \\ M \xrightarrow{d_1} \varnothing, \quad R \xrightarrow{d_2} \varnothing, \quad C_1 \xrightarrow{a_1} R, \quad C_1 \xrightarrow{a_2} M, \quad C_2 \xrightarrow{b_1} 2R, \quad C_2 \xrightarrow{b_2} C_1. \end{gathered}$$

For convenience, let $m$ denote the number of mRNA, let $r$ denote the number of microRNA, let $n_i$ denote the number of complex $C_i$.

We next use Holimap to map the gene network to the following linear network:

$$G_1 \xrightarrow{\sigma_{b1}} G_1^*, \quad G_1^* \xrightarrow{\sigma_{u1}} G_1,$$
$$G_1 \xrightarrow{\rho_{u1}} G_1 + M, \quad G^* \xrightarrow{\rho_{b1}} G_1^* + M, \quad M \xrightarrow{\tilde{d}_1} \varnothing.$$

Similarly to the derivation in previous sections, the effective parameter $\tilde{d}$ should be determined by

$$\tilde{d}_1\langle m\rangle = d_1\langle m\rangle + \alpha\langle mr\rangle - (\beta + a_2)\langle n_1\rangle. \tag{34}$$

In summary, once we have computed the values of moments, we can use Eq. (34) to determine the values of the effective parameter.

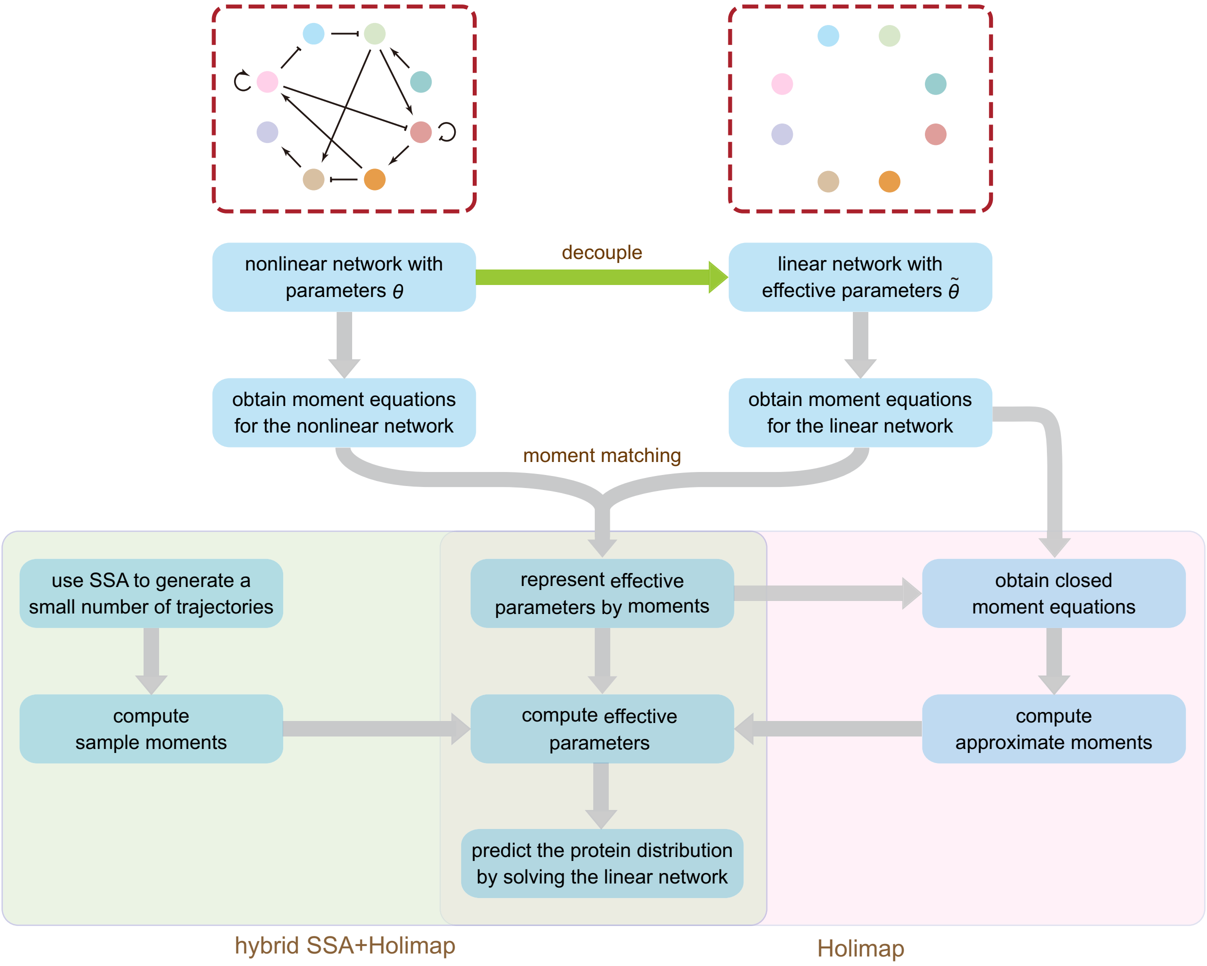


Figure 1: **Flow chart of the Holimap algorithm for a general gene network.** The pink frame shows the algorithm for the standard Holimap, and the green frame shows the algorithm for the hybrid approach SSA and Holimap.

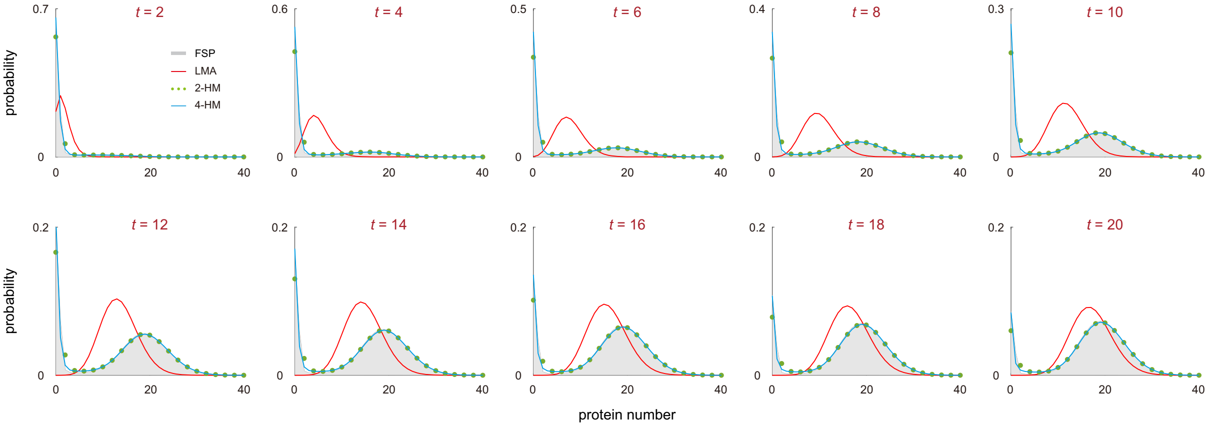


Figure 2: **Holimaps for autoregulatory gene circuits.** Comparison of the time-dependent protein distributions computed using the FSP, LMA, 2-HM, and 4-HM. The parameters are chosen as $h = 2, d = 1, B = 0.1, \rho_u = 5, \rho_b = 200, \sigma_u = 100, \sigma_b = 20$.

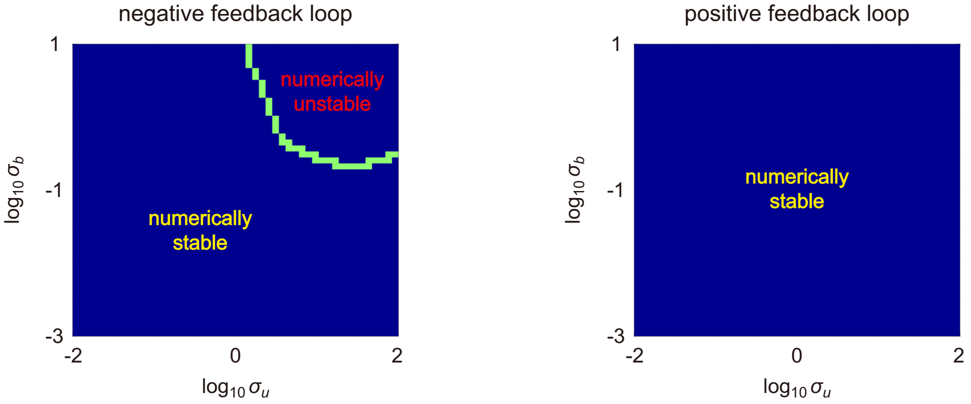


Figure 3: **Numerical stability for the 4-HM in autoregulatory feedback loops.** The left (right) panel shows the numerically stable parameter region and the numerically unstable parameter region for the 4-HM in negative (positive) feedback loops at time $t = 1$. The green curve separates the numerically stable and unstable regions in the $\sigma_u$-$\sigma_b$ plane. Clearly the 4-HM may lead to numerical instability when the binding and unbinding rates, $\sigma_b$ and $\sigma_u$ are large. However, for positive feedback loops, we do not observe numerical instability for the 4-HM. The parameters are chosen as $h = 2, d = 1, B = 0.1$; the protein burst frequencies are chosen as $\rho_u = 200$, $\rho_b = 1$ for the negative feedback loop and $\rho_u = 1$, $\rho_b = 200$ for positive feedback loops.

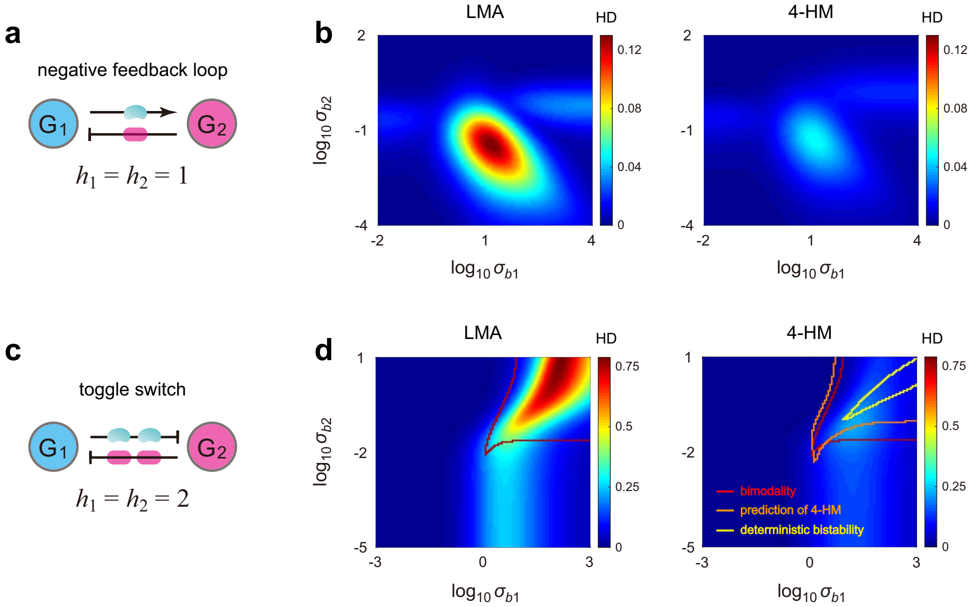


Figure 4: **Holimaps for two-node gene networks.** (**a**) A negative feedback loop with no cooperative binding. (**b**) Heat plots of the HDs for the LMA and 4-HM as functions of the binding rates $\sigma_{b1}$ and $\sigma_{b2}$. Here the HD represents the Hellinger distance between the real and approximate steady-state distributions of protein $P_2$ numbers. The parameters are chosen as $h_1 = h_2 = 1, d_1 = d_2 = 1, \rho_{u1} = 21, \rho_{b1} = 7, \rho_{u2} = 0, \rho_{b2} = 3, \sigma_{u1} = \sigma_{u2} = 5$. (**c**) A toggle switch with cooperative binding. (**d**) Same as (b) but for the toggle switch. The red curve encloses the true bimodal region computed using FSP, the orange curve encloses the bimodal region predicted by the corresponding LMA, and the yellow curve encloses the region of deterministic bistability. The parameters are chosen as $h_1 = h_2 = 2, d_1 = d_2 = 1, \rho_{u1} = 18, \rho_{b1} = 1, \rho_{u2} = 2, \rho_{b2} = 0, \sigma_{u1} = \sigma_{u2} = 5$.

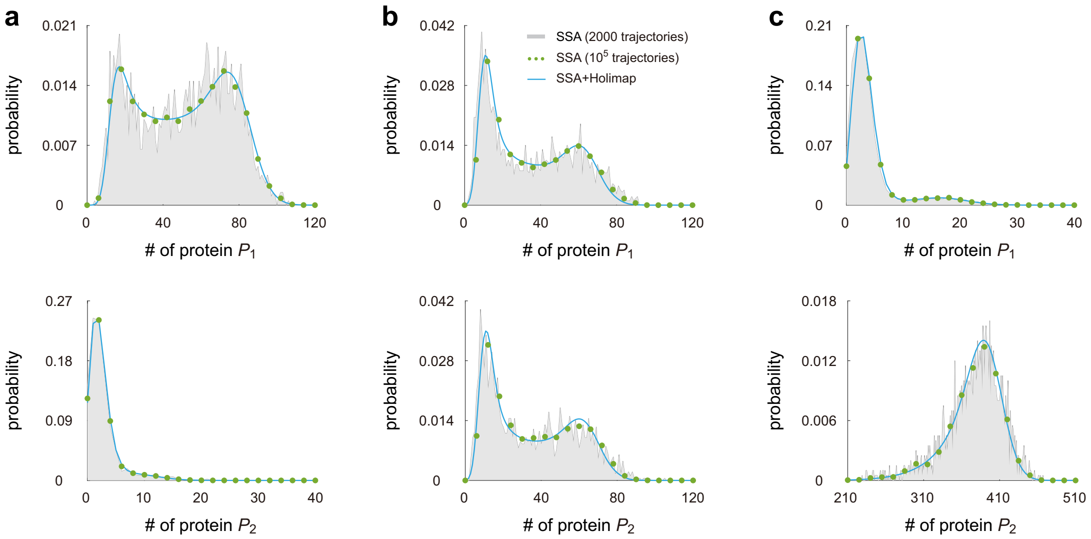


Figure 5: **Holimap for the post-translational network shown in Fig. 6(a) in the main text.** (**a**) Protein number distributions when protein $P_2$ is very scarce compared to protein $P_1$. (**b**) Protein number distributions when proteins $P_1$ and $P_2$ interact at comparable concentrations. (**c**) Protein number distributions when protein $P_2$ is very abundant compared to protein $P_1$. The parameters are chosen as $h_1 = h_2 = 1, d_1 = d_2 = 1, \alpha = 0.3, \rho_{u1} = 14, \rho_{b1} = 84, \sigma_{u1} = \sigma_{u2} = 0.8, \sigma_{b1} = \sigma_{b2} = 0.029$. The remaining parameters are chosen as $\rho_{u2} = 0.2\rho_{u1}, \rho_{b2} = 0.2\rho_{b1}$ for (a), $\rho_{u2} = \rho_{u1}, \rho_{b2} = \rho_{b1}$ for (b), and $\rho_{u2} = 5\rho_{u1}, \rho_{b2} = 5\rho_{b1}$ for (c).

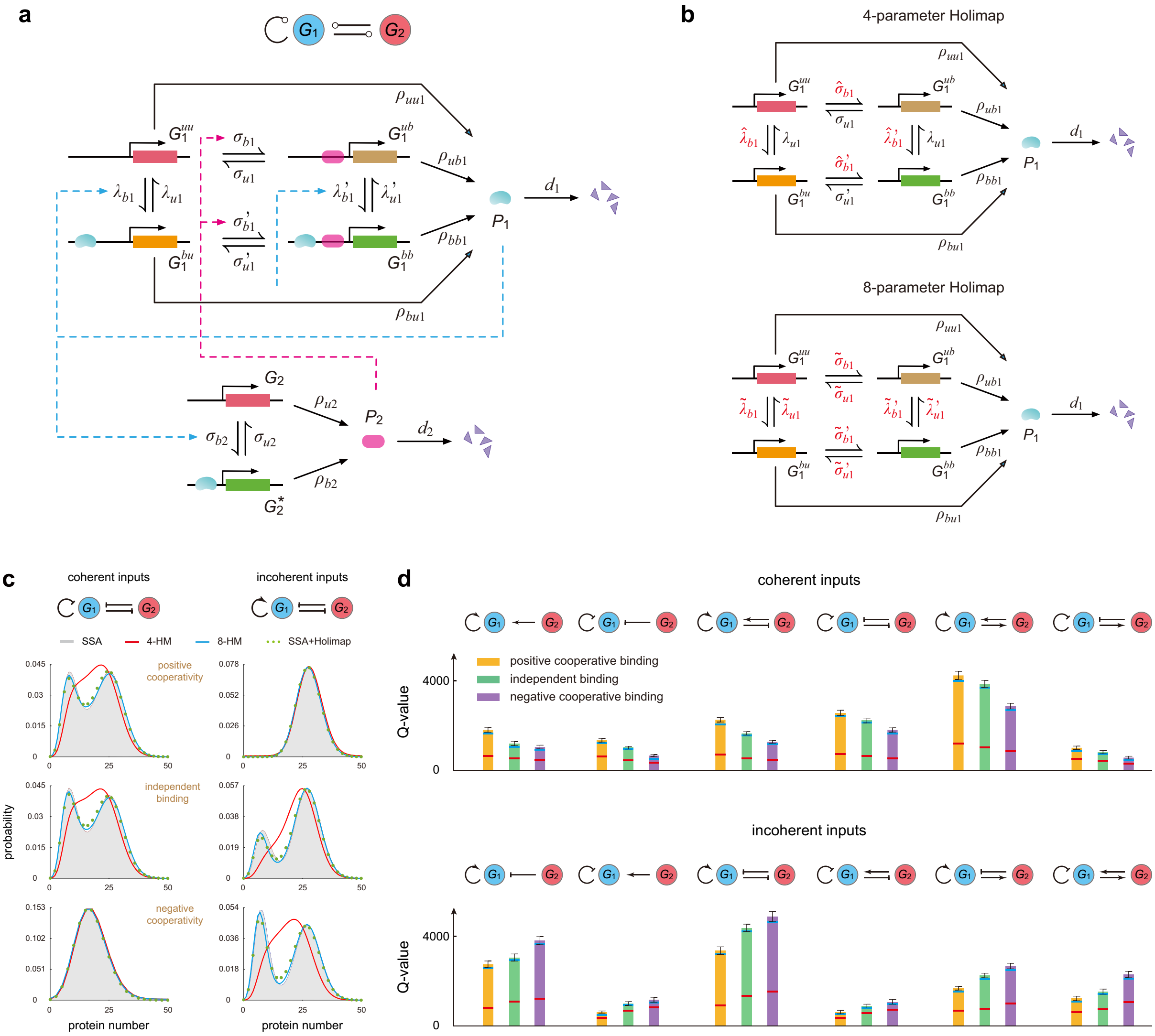

Figure 6: **Two-node gene networks with transcription factor interactions.** (**a**) Stochastic model of a two-node regulatory network, where gene $G_2$ is regulated by gene $G_1$ and gene $G_1$ is regulated by both gene $G_2$ and itself. Feedback is mediated by binding of protein $P_1$ to gene $G_2$ and cooperative binding of proteins $P_1$ and $P_2$ to gene $G_1$. Specifically, the promoter of gene $G_1$ has two binding sites, one for protein $P_1$ and the other for protein $P_2$. According to whether the two binding sites are occupied or not, there are a total of four states for gene $G_1$ with four binding rates and four unbinding rates. The binding of the two proteins may have three types of interactions: independent binding, positive cooperative binding, and negative cooperative binding. (**b**) Illustration of the 4-HM and 8-HM applied to predict the distribution of protein $P_1$ numbers of the nonlinear network. In the 4-HM, the four binding rates associated with gene $G_1$ are replaced by four effective parameters in the linear network. In the 8-HM, all the binding and unbinding rates are modified. (**c**) Steady-state distribution of protein $P_1$ numbers for two types of network topologies (networks with coherent and incoherent inputs) and three types of transcription factor interactions: positive cooperativity (upper), independent binding (middle), and negative cooperativity (lower). The protein distributions are computed using the SSA, 4-HM, 8-HM, and SSA+8-HM. Here coherent (incoherent) inputs mean that the autoregulation of gene $G_1$ and its regulation by gene $G_2$ have the same (opposite) signs. (**d**) $Q$-values for all 12 possible network topologies and for three types of transcription factor interactions. For each topology and each type of interaction, we randomly select $10^4$ sets of model parameters and compute the number of sets $Q$ that yield a bimodal steady-state distribution using FSP (solid bars), 4-HM (red line) and 8-HM (blue line). The error bar shows the standard deviation of the FSP prediction for three independent batches of $10^4$ parameters. Here we apply FSP rather than SSA because the latter often results in a non-smooth distribution, from which we may incorrectly identify the number of modes. See Supplementary Note 1 for the technical details of this figure.
